# COMPENSATION FOR CHRONIC OXIDATIVE STRESS IN ALADIN NULL MICE

**DOI:** 10.1101/164442

**Authors:** Ramona Jühlen, Mirko Peitzsch, Sebastian Gärtner, Dana Landgraf, Graeme Eisenhofer, Angela Huebner, Katrin Koehler

## Abstract

**Background:** Mutations in the *AAAS* gene coding for the nuclear pore complex protein ALADIN lead to the autosomal recessive disorder triple A syndrome. Triple A patients present with a characteristic phenotype including alacrima, achalasia and adrenal insufficiency. Patient fibroblasts show increased levels of oxidative stress and several *in vitro* studies demonstrated that the nucleoporin ALADIN is involved in the cellular oxidative stress response and in adrenal steroidogenesis. We showed that ALADIN knock-out mice lack a phenotype resembling human triple A syndrome. Thus, we hypothesized that application of chronic oxidative stress by ingestion of paraquat will generate triple A-like phenotype in ALADIN null mice.

**Results:** We demonstrate that ALADIN knock-out mice present with an unexpected compensated glutathione metabolism still lacking a phenotype resembling human triple A syndrome after application of chronic oxidative stress. We could not observe increased levels of oxidative stress and alterations in adrenal steroidogenesis in mice depleted for ALADIN.

**Conclusions:** This study stresses the species-specific role of the nucleoporin ALADIN presenting a novel compensatory mechanism of the cellular glutathione redox response and shedding light on the role of ALADIN in the cell.

## BACKGROUND

The triple A syndrome (OMIM #231550), a rare autosomal recessive disorder, is caused by homozygous or compound heterozygous mutations in the *AAAS* (**a**chalasia-**a**drenal insufficiency-**a**lacrima **s**yndrome) gene encoding the nucleoporin ALADIN (**al**acrima-**a**chalasia-a**d**renal **i**nsufficiency **n**eurologic disorder) [1,2]. Triple A patients present with the characteristic triad of adrenocorticotropic hormone-resistant adrenal insufficiency, achalasia of the stomach cardia and alacrima in combination with progressive neurological impairments [3]. Phenotypic appearance of all symptoms is heterogeneous and highly variable. Adrenal atrophy may occur later in life and may develop gradually [4,5]. ALADIN is anchored within the nuclear pore complex by the transmembrane nucleoporin NDC1 (nuclear division cycle 1 homologue (*S. cerevisiae*)) [6,7]. Rabut et al. suggested that ALADIN forms part of the structural backbone of the nuclear pore complex but is not needed itself for integrity of the complex [8].

In contrast to other organs with high metabolic rates the adrenal gland has high levels of enzymatic and non-enzymatic anti-oxidants [9]. Imbalances in reactive oxygen species (ROS) result in cellular oxidative stress and have been implicated in a variety of diseases [9]. Adrenocortical mitochondrial steroidogenesis significantly adds to ROS formation in the cell because uncoupling of the cytochrome P450 enzyme (CYP) redox reaction can occur in several steps of the reaction [10,11]. Under these circumstances superoxide anions and hydrogen peroxide can leak and escape from the redox reaction [10]. Therefore, an equilibrated level of anti-oxidative mechanisms is of high importance in adrenocortical cells.

A sound body of published work has reported that ALADIN is involved in the cellular oxidative stress response *in vitro* in adrenocortical and fibroblast cells but the role of ALADIN in adrenal steroidogenesis and how this might contribute to adrenal atrophy in triple A patients remains largely unknown [12–17]. We have shown that depletion of ALADIN in human adrenocortical carcinoma cells leads to an alteration in glucocorticoid and androgen steroidogenesis [13]. Recently, we identified progesterone receptor membrane compartment 2 (PGRMC2) as a novel protein interactor of ALADIN [18]. Microsomal PGRMC2 itself seems to be involved in adrenal steroidogenesis either by regulating heme synthesis, the prosthetic group of microsomal CYPs, or by acting as an electron donor for several CYPs [19,20].

Here, we sought to verify the critical role for ALADIN in the cellular redox regulation in ALADIN null mice. Female homozygous mice deficient for *Aaas* are sterile/infertile but otherwise ALADIN null mice present with a mild phenotype [21]. Carvalhal et al. postulated that female sterility in ALADIN-deficient mice is caused by impaired chromosomal segregation and maturation of oocytes [22]. Most recently, it was shown that conditional ablation of ALADIN interactor PGRMC2 from the female reproductive tract results in reproductive senescence [23].

We hypothesized that application of oxidative stress using paraquat in mice deficient for ALADIN will generate symptoms seen in triple A patients which the animals lack under basal conditions. In order to increase the sensitivity for oxidative stress we used, besides our *Aaas* knock-out (KO) mice, offspring from intercrossed heterozygous (Het) *Sod2* female mice and *Aaas* KO male mice to obtain *Aaas* KO/*Sod2* Het mice.

## MATERIALS AND METHODS

### •Experimental animals and treatments

All mice were housed in the animal care facility (Experimental Center) of the Technical University Dresden, Dresden, Germany. All procedures were approved by the Regional Board for Veterinarian Affairs, Dresden, Germany (AZ 24-9168.11-1/-2010-49) in accordance with the institutional guidelines for the care and use of laboratory animals. Animals were group housed except during actual experimental procedures, when single housing was required. Mice were kept under specific-pathogen-free conditions at a constant temperature (22 ± 1 °C) and a 12 hours light-dark cycle at all times. Mice were weaned onto ssniff R/M-H (19% protein, 4.9% fibers, 3.3% fat, 12.2 MJ/kg). (ssniff GmbH, Soest, Germany) if not stated otherwise and drank water ad libitum. *Aaas-*deficient mice were generated as described previously [21] and backcrossed to strain C75BL/6J for ten generations. A heterozygous *Sod2* mouse strain was obtained from The Jackson Laboratory, Bar Harbor, ME USA (Strain #002973 B6.129S7-Sod2tm1Leb/J). Heterozygous *Sod2* female mice were intercrossed for two generations with *Aaas* KO male mice to obtain *Aaas* KO/*Sod2* Het mice.

For chronic oxidative stress exposure one-year-old adult male mice of three different genotypes [wild-type (WT) (n=16), *Aaas* KO (n=16) and *Aaas* KO/*Sod2* Het (n=10)] were used and randomly divided into two groups (stress and control group). All were placed on a commercial diet (ssniff R/M-H) for 3 days to allow acclimation to these conditions. Mice were then fed with paraquat diet (0.25 g/kg diet) (ssniff GmbH) in the stress group and with control diet (ssniff GmbH) for 11 days. Body weight and diet weight were determined every day during the feeding period. At the end (day 11) of the feeding period animals were sacrificed. Lung and liver were surgically removed, washed in ice-cold PBS and weighed. Different parts of the liver were prepared for glutathione measurement and assessment of lipid peroxidation. Adrenals and liver sections were surgically excised and quickly frozen in liquid nitrogen and stored at -80 °C before RNA extraction.

### •Hepatic glutathione assay

Small samples (40-100 mg) of liver tissue were rapidly cut on an ice-cold petri dish ensuring to prevent oxidation of reduced glutathione (GSH) to oxidized glutathione (GSSG) during preparation. Each small sample was immediately placed with a forceps in liquid nitrogen. Samples in the tubes were re-weighed and the weight of the tissue was determined. Ten volumes of ice-cold 5% sulfosalicylic acid (Carl Roth, Karlsruhe, Germany) were added to each tube, the sample was transferred to a tissue grinder and homogenized until evenly suspended. The suspension was added to the same tube and centrifuged at 4 °C at 14000x g for 10 minutes. The supernatant was transferred to a new tube and equal volume of ice-cold 500 mM HEPES (pH 8) (Gibco, Thermo Fisher Scientific, Schwerte, Germany) were added.

Each sample was diluted 60-fold in ice-cold 250 mM HEPES (pH 7.5) to be in linear detection range for measurement of total and oxidized glutathione using the GSH/GSSG-Glo assay (Promega, Mannheim, Germany). Measurements were done in duplicate as outlined in the protocol of the manufacturer and as reported elsewhere [13].

### •Hepatic lipid peroxidation measurement

End-products of hepatic lipid peroxidation, malondialdehyde precursors and other thiobarbituric acid reactive substances (TBARS), were extracted from liver sections as described before by centrifugation at 1600x g for 10 minutes [24]. TBARS were quantified in triplicate spectrophotometrically at 535 and 520 nm as outlined previously [24] on a 96-well culture dish (200 μl/well) (Corning Costar, Kaiserslautern, Germany) using a Infinite 200 PRO Microplate Reader with the Magellan Data Analysis Software v6.6 (Tecan Group AG, Männedorf, Switzerland).

### •RNA extraction, cDNA synthesis and quantitative real-time PCR using TaqMan

Total RNA from frozen murine liver and adrenals was isolated, purity assessed, reverse transcribed and qPCR amplifications in 20 μl total volumes performed as outlined elsewhere [18].

As reference gene for normalization beta-actin was evaluated and used. Positive controls contained a random mix of cDNA and negative controls contained nuclease-free water instead of cDNA. In all real-time qPCR experiments relative gene expression was calculated by the C_t_ method using standard and semi-log plots of amplification curves. In all results repeatability was assessed by standard deviation of triplicate C_t_ s and reproducibility was verified by normalizing all real-time RT-PCR experiments by the C_t_ of each positive control per run.

Primers for the amplification of the target sequence of beta actin (*Actb*), *Cyp11a1, Cyp11b1,Cyp11b2, Cyp21a1*, glutathione peroxidase 1 (*Gpx*1), glutathione reductase (*Gsr*), heme oxygenase 1 (*Hmox1*), hydroxy-delta-5-steroid dehydrogenase (*Hsd3b2*), nicotinamide nucleotide transhydrogenase (*Nnt*), superoxide dismutase 2 (*Sod2*) and steroidogenic acute regulatory protein (*Star*) were designed using Primer Express 3.0 (Applied Biosystems, Life Technologies, Darmstadt, Germany) and compared to the murine genome database for unique binding using BLAST search (https://blast.ncbi.nlm.nih.gov/Blast.cgi). The primer sequences and gene accession numbers are listed in **Table S1**.

The guidelines of the Minimum Information for Publication of Quantitative Real-Time PCR Experiments were followed in this study to allow more reliable interpretation of real-time RT-PCR results [25].

### •LC-MS/MS measurement of steroids

Blood for plasma steroid measurement by liquid chromatography tandem mass spectrometry (LC/MS-MS) was collected by cardiac puncture. Plasma steroids pregnenolone (Preg), progesterone (P), 17-hydroxyprogesterone (17OHP), deoxycorticosterone (DOC), corticosterone (B), aldosterone (ALDO), androstenedione (AE), dehydroepiandrosterone (DHEA) and dehydroepiandrosterone sulfate (DHEAS) were determined simultaneously by LC-MS/MS as reported previously [26]. Quantification of steroid levels was done by comparisons of ratios of analyte peak area obtained from plasma samples to the respective peak area of stable isotope labelled internal standard calibrators.

### •Histology

Sections of brain, duodenum, liver and lung were washed in PBS and fixed in 4% formaldehyde (SAV LP, Flinsbach, Germany) for 24 hrs. Organs were then transferred to PBS and prepared for histology at the Histology Facility of the Joint Technology Platform (Technische Universität Dresden, Biotec, CRTD).

Tissues were embedded into paraffin with the Microm STP 420 D dehydration/infiltration unit (Thermo Fisher Scientific, Waltham, Massachusetts, USA) and the EGF 1160 embedding station (Leica, Wetzlar, Germany). This included stepwise dehydration in a graded alcohol series, transfer to xylol, as well as paraffin infiltration and sample orientation. Paraffin-embedded samples were sectioned using a Microm HM 340E (Thermo Fisher Scientific) and stained with hematoxylin-eosin (Carl Roth).

### •Statistical analysis

Statistical analyses were made using the open-source software R version 3.3.2 and R Studio version 1.0.136 (R Core Team, 2016). Unpaired Wilcoxon–Mann–Whitney U-test was performed. During evaluation of the results a confidence interval alpha of 95% and P-values lower than 0.05 were considered as statistically significant.

## RESULTS

### •Chronic oxidative stress is not detected by increased *Hmox1* expression

Assessment of the level of oxidative stress was done by measuring adrenal Hmox1 gene expression by qPCR. *Hmox1* is a widely-used redox-regulated gene whose transcriptional activation is dependent on upstream transcriptional regulators which are induced by a broad spectrum of conditions involving oxidative stress, nitrosative stress, thiol-reactive substances and cytokines [27]. We could not see an increased expression of *Hmox1* in animals under paraquat diet compared to control diet (**Fig. 1A**). However, under control diet the expression was significantly decreased in *Aaas* KO/*Sod2* Het compared to *Aaas* KO animals.

**Figure 1.**
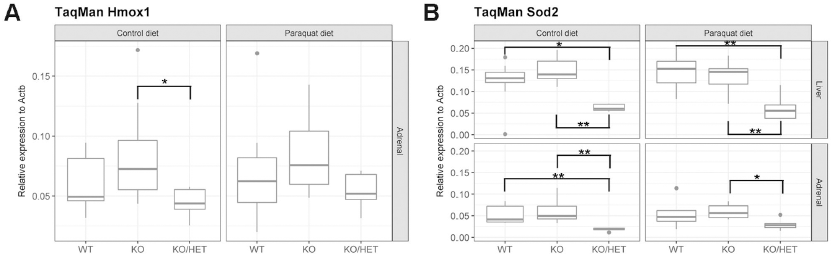
Expression analysis of (A) redox-regulated adrenal *Hmox1* and (B) adrenal and hepatic *Sod2*. Mice were fed with paraquat diet (0.25 g/kg diet) and with control diet for 11 days. P-values: *P<0.05, **P<0.01. Significant differences were measured with unpaired Wilcoxon– Mann–Whitney U-test. Boxplot widths are proportional to the square root of the samples sizes. Whiskers indicate the range outside 1.5 times the inter-quartile range above the upper and below the lower quartile. Outliers were plotted as dots.

Hepatic and adrenal *Sod2* expression was about two-fold diminished in *Aaas* KO/*Sod2* Het mice under control or paraquat diet compared to WT and *Aaas* KO mice of the same diet (**Fig. 1B**).

### •Adrenal steroid output is not affected by chronic oxidative stress exposure

The expression of *Star* was increased in *Aaas* KO versus WT animals after paraquat diet (**Fig. 2A**). Furthermore, *Aaas* KO/*Sod2* Het mice under paraquat diet presented with decreased expression of *Star* compared to *Aaas* KO mice of the same diet but neither expression levels of *Cyp21a1, Cyp11a1, Cyp11b1, Cyp11b2*, and *Hsd3b2* were changed nor could we see a specific effect depending on genotype of the mice (**Fig. S1A-E**).

**Figure 2.**
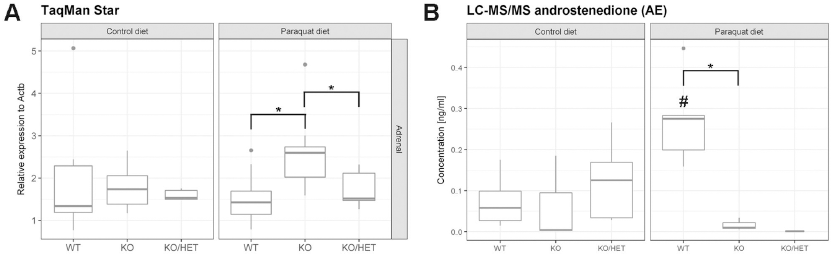
Oxidative stress affects (A) expression of *Star* and (B) testicular synthesis androstenedione. Mice were fed with paraquat diet (0.25 g/kg diet) in the stress group and with control diet for 11 days. P-values:* P<0.05. Significant differences were measured with unpaired Wilcoxon–Mann–Whitney U-test. Boxplot widths are proportional to the square root of the samples sizes. Whiskers indicate the range outside 1.5 times the inter-quartile range above the upper and below the lower quartile. Outliers were plotted as dots.

Plasma levels of Preg, P, DOC, B, ALDO and DHEAS were not significantly altered upon paraquat diet or in between the different genotypes (**Fig. S2A-F**). Plasma levels of 17OHP and DHEA were under detection threshold. Production of AE which in mice is only synthesized in gonads was about five-fold increased in WT animals after ingestion of paraquat compared to WT animals of the control diet (**Fig. 2B**). Furthermore, AE levels in *Aaas* KO versus WT animals were about 25-fold decreased after paraquat diet (**Fig. 2B**).

### •Paraquat diet and ALADIN depletion decrease body weight gain

In the control diet food intake over 11 days of experimental procedure was significantly decreased in *Aaas* KO/*Sod2* Het mice compared to WT mice (**Fig. 3A**). We also saw a lowered food intake in *Aaas* KO animals but it appeared not to be significant. Weight gain in *Aaas* KO and *Aaas* KO/*Sod2* Het mice was about two-fold diminished in the control diet compared to the WT (**Fig. 3B**).

**Figure 3.**
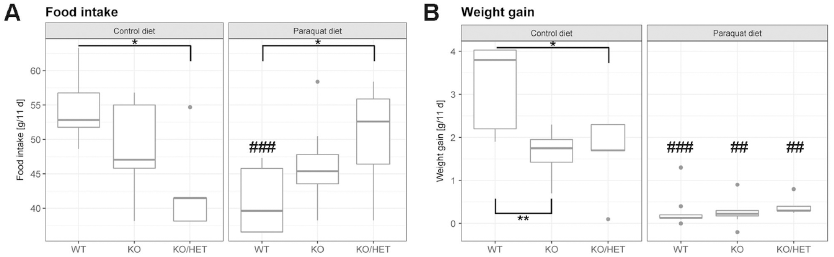
Alteration of (A) food intake and (B) body weight gain by oxidative stress. Mice were fed with paraquat diet (0.25 g/kg diet) in the stress group and with control diet for 11 days. Body and diet weight were determined every day during the feeding period. P-values: (between different genotypes in one diet) *P<0.05, **P<0.01 and (between different diets in one genotype) ## P<0.01, ### P<0.001. Significant differences were measured with unpaired Wilcoxon–Mann– Whitney U-test. Boxplot widths are proportional to the square root of the samples sizes. Whiskers indicate the range outside 1.5 times the inter-quartile range above the upper and below the lower quartile. Outliers were plotted as dots.

WT mice on the paraquat diet fed less compared to the control diet; however, food intake in *Aaas* KO/*Sod2* Het mice was higher versus the WT (**Fig. 3A**). Accordingly, WT animals gained about 20-fold less weight compared to the control diet and weight gain was also about four-fold lowered in *Aaas* KO and *Aaas* KO/*Sod2* Het animals versus the control diet despite increased food intake (**Fig. 3A-B**).

### •Hepatic glutathione levels are balanced in ALADIN null mice

Hepatic GSH/GSSG ratios in *Aaas* KO/*Sod2* Het animals either under control or paraquat diet were about five-fold increased compared to WT animals under the same diet (**Fig. 4A**). Additionally, paraquat diet increased the ratio significantly in *Aaas* KO/*Sod2* Het animals compared to the same genotype under control diet. GSH/GSSG ratios of *Aaas* KO mice rather presented like WT mice.

**Figure 4.**
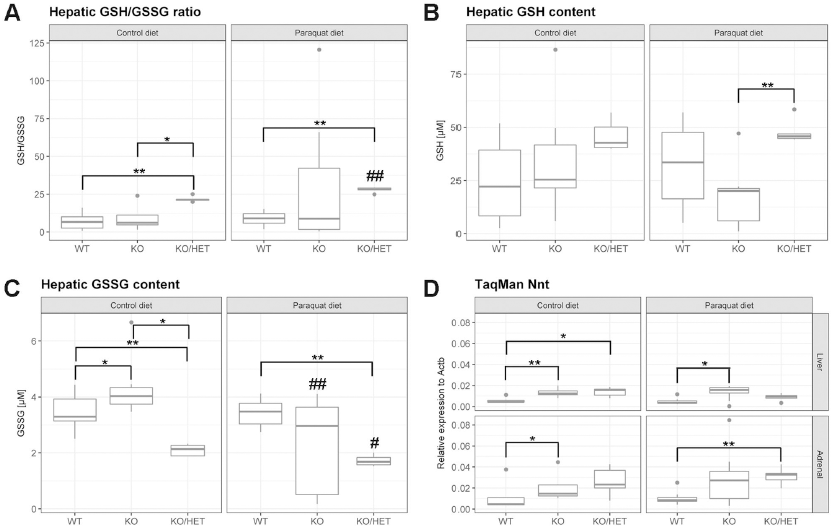
Balance of hepatic glutathione levels in ALADIN null mice. Mice were fed with paraquat diet (0.25 g/kg diet) in the stress group and with control diet for 11 days. GSH, reduced glutathione. GSSG, oxidized glutathione. P-values: (between different genotypes in one diet) *P<0.05, **P<0.01 and (between different diets in one genotype) # P<0.05, ## P<0.01. Significant differences were measured with unpaired Wilcoxon–Mann–Whitney U-test. Boxplot widths are proportional to the square root of the samples sizes. Whiskers indicate the range outside 1.5 times the inter-quartile range above the upper and below the lower quartile. Outliers were plotted as dots.

GSH concentrations were higher in *Aaas* KO/*Sod2* Het livers either under control or paraquat diet versus WT and *Aaas* KO mice under the same diet (**Fig. 4B**). Hepatic GSH content in *Aaas* KO mice were comparable to WT mice.

Similarly, hepatic GSSG concentrations were about two-fold diminished in *Aaas* KO/*Sod2* Het animals compared to WT and *Aaas* KO animals under the same diet (**Fig. 4C**). Furthermore, GSSG concentrations decreased in *Aaas* KO/*Sod2* Het mice when under paraquat diet. Interestingly, GSSG content in *Aaas* KO mice under control diet was significantly higher compared to WT and *Aaas* KO/*Sod2* Het mice. This effect was reversed under paraquat diet: GSSG concentration in *Aaas* KO animals decreased compared to control diet.

We could see no alteration in the expression of *Gpx1* in control or paraquat diet animals or in the different genotypes (**Fig. S3A**). However, *Aaas* KO/*Sod2* Het mice under paraquat diet presented with decreased expression of *Gsr* compared to *Aaas* KO mice (**Fig. S3B**). Most strikingly, hepatic and adrenal expression of *Nnt* was about two-fold increased in *Aaas* KO and *Aaas* KO/*Sod2* Het mice under control diet versus the WT (**Fig. 4D**). Under paraquat diet hepatic *Nnt* expression was still higher in *Aaas* KO animals compared to WT animals and adrenal *Nnt* expression was significantly increased in *Aaas* KO/*Sod2* Het mice under paraquat diet.

Relative liver and lung weights and hepatic TBARS values were not altered upon oxidative stress exposure using a paraquat diet or in the different genotypes (**Figs. S4 and S5**). No pathological differences in histology sections of brain, duodenum, liver and lung could be found (data not shown).

## DISCUSSION

In the present study we investigated the role of the nucleoporin ALADIN in chronic paraquat-induced oxidative stress in male mice. ALADIN-deficient mice lack a triple A syndrome-characteristic phenotype [21]. Previous studies have demonstrated that ALADIN employs a crucial role in the redox response of the cell *in vitro* [12–17]. Triple A patients as well suffer from increased cellular oxidative stress as preliminary shown by Fragoso et al. [28]. Thus, we hypothesized that chronic oxidative stress will unmask the distinct phenotype in ALADIN null mice.

Overall, after chronic oxidative stress exposure we did not see a triple A syndrome-characteristic phenotype in mice depleted for ALADIN. Previous to our study a pilot experiment using acute oxidative stress by injection with paraquat i.p. (25 mg/kg body weight) in mice was performed but no involvement of ALADIN in the acute oxidative stress response was obtained (data not shown). Our murine *in vivo* result in this study is contrary to various human *in vitro* cell systems in which upon depletion of ALADIN a disturbed redox homeostasis and altered adrenal steroidogenesis were seen [12–14]. We assume that this discrepancy is either a result of the species-specific role of ALADIN or of the experimental nature of the study comparing *in vitro* with *in vivo* models.

In more detail, our data indicate that on the one hand, mice depleted for ALADIN during basal conditions and after chronic oxidative stress exposure sustain with balanced hepatic glutathione levels by up-regulation of *Nnt* resulting in a WT-like phenotype. On the other hand, *Aaas* KO/*Sod2* Het mice under basal conditions increase hepatic glutathione levels by increasing *Nnt* expression. This effect was intensified after chronic oxidative stress exposure. In the cell transmembrane nicotinamide nucleotide transhydrogenase (NNT) plays a key role in the mitochondrial defense system against reactive oxygen species (ROS) by producing NADPH (**Fig. 5**). NADPH is in turn consumed by glutathione reductase (GSR) maintaining reduced glutathione (GSH) levels from oxidized glutathione (GSSG) [29]. ROS, i.e. superoxide anions, leaking during mitochondrial aerobic respiration or produced by exogenous stressors are converted to hydrogen peroxide by mitochondrial superoxide dismutase (SOD2). Hydrogen peroxide is then neutralized to water-consuming GSH by several peroxidases (GPX).

**Figure 5.**
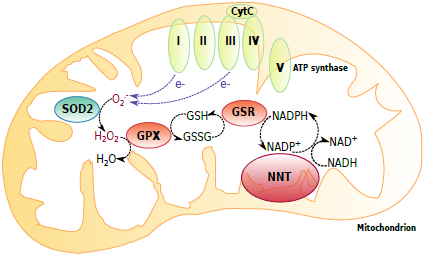
Mitochondrial redox defense system. Transmembrane nicotinamide nucleotide transhydrogenase (NNT) contributes to the mitochondrial redox defense system by producing NADPH. NADPH is consumed by glutathione reductase (GSR) maintaining reduced glutathione (GSH) levels from oxidized glutathione (GSSG). Electrons leaking during mitochondrial aerobic respiration result in superoxide anion radicals (O_2_^.-^) and are converted to hydrogen peroxide (H_2_O_2_) by mitochondrial superoxide dismutase (SOD2). Hydrogen peroxide is neutralized to water (H_2_O) consuming GSH by glutathione peroxidase (GPX).

It has already been shown that heterozygous deficiency for *Sod2* in mice activates mitochondrial uncoupling to reduce ROS production and increases aerobic glycolysis by a free radical-mediated mechanism [30]. Mice heterozygous deficient for *Sod2* exhibit increased levels of ROS and shift from mitochondrial oxidative phosphorylation to a cytosolic glycolytic pathway [30]. During aerobic glycolysis a high rate of energy is produced by metabolizing glucose into pyruvate which then feeds into cytosolic lactic acid fermentation rather than mitochondrial oxidation, commonly known as the Warburg Effect [30,31]. In the present study it can thus be assumed that the phenotypic effects seen in *Aaas* KO/*Sod2* Het mice are caused by both the Warburg Effect and increased expression of *Nnt.* This additive effect in turn leads to transient increase of glutathione oxidative capacity and to an enhancement of the compensatory effect seen in *Aaas* KO mice which lack typical symptoms of triple A syndrome. Thus, we present that ALADIN plays a crucial role in regulating NADPH levels in the cell and concomitantly enhances oxidative capacity of glutathione by altered gene expression of NNT. Gene down-regulation of *Nnt* has been associated with age-related neurodegeneration in Alzheimer disease-like mouse neurons [32]. It has been reported that NAD(P)H redox control is more critical than GSH content in promoting neurodegeneration [32]. This result partly explains why mice depleted for ALADIN do not present with a triple A syndrome-distinct phenotype but rather behave like WT animals. In view of its cellular localization at the nuclear pore we hypothesize that ALADIN plays a role in regulating the export of *Nnt* mRNA through the nuclear pore complex and thus, in balancing levels of NADPH.

We based our study of chronic paraquat-induced oxidative stress on the work of Aoki and colleagues in which a 0.025% paraquat enriched diet was also used to induce oxidative stress in four-week-old juvenile male rats [33]. In contrast to our results Aoki et al. found that by feeding rats the paraquat diet animals suffered from elevated hepatic lipid (TBARS) and glutathione (GSSG) oxidation, liver organ shrinkage and lung enlargement [33]. We could not reproduce these results in our mice. This may be due to the different age of the animals or to different anti-oxidant defenses in the two rodent species. Results from Aoki et al. regarding food intake and body weight gain were consistent to our study [33]. Here, we show that depletion of ALADIN in mice negatively affected body weight gain under normal control and paraquat diet. This result is underlined by increased food intake under paraquat diet in these animals.

## CONCLUSIONS

Our *in vivo* study is the first to highlight a species-specific role of the nucleoporin ALADIN. Our study implies a complex cellular system involved to compensate a depletion of ALADIN which seems to have an important task in balancing NADPH levels in the cell. Future research shall address which other and how these pathways are involved in a possible compensating mechanism clarifying the role of ALADIN in the pathogenesis in triple A syndrome.

## DECLARATIONS

### Availability of data and material

The datasets analyzed during the current study are available from the corresponding author on reasonable request. All data generated or analyzed during this study are included in this published article and its supplementary information files.

## Author's contributions

RJ, SG, AH and KK conceived and designed the experiments. RJ, MP, SG, DL and KK performed all experiments. RJ, MP, KK and AH analyzed the data. RJ wrote the paper. MP, SG, GE, AH and KK assisted with improving the manuscript. All authors read the final version of the manuscript and gave their permission for publication.

### Acknowledgements

We thank Michael Haase for analyzing murine histology sections.

